# No evidence for immune-gene specific signals of selection in termites

**DOI:** 10.1101/783738

**Authors:** Karen Meusemann, Judith Korb, Maximilian Schughart, Fabian Staubach

## Abstract

It has been hypothesized that selection pressure from pathogens plays an important role in shaping social evolution. Social behaviour, in particular brood care, is associated with pathogen pressure in wood-dwelling “lower” termites. Yet, generally pathogen pressure is low in wood-dwelling termite species that never leave the nest except for the mating flight. In comparison, pathogen pressure is higher in species that leave the nest to forage, and thus constantly encounter a diversity of microbes from their environment. We hypothesized that such differences in pathogen pressure are also reflected by differences in the intensity of natural selection on immune genes. We tested this hypothesis in a phylogenetic framework, analyzing rates of non-synonymous and synonymous substitutions on single-copy immune genes. Therefore, we leveraged recent genomic and transcriptomic data from eight termite species, representing wood-dwelling and foraging species as well as 14 additional species spanning the winged insects (Pterygota). Our results provide no evidence for a role of pathogen pressure in selection intensity on single-copy immune genes. Instead, we found evidence for a genome-wide pattern of relaxed selection in termites.

**Life Science Identifiers (as available Zoobank):** *Ephemera danica*:

urn:lsid:zoobank.org:act:06633F75-4809-4BB3-BDCB-6270795368D5

*Coptotermes* sp.

urn:lsid:zoobank.org:pub:D6724B7F-F27A-47DC-A4FC-12859ECA0C71

*Blattella germanica*:

rn:lsid:zoobank.org:pub:1EA126BA-E9D2-4AA6-8202-26BA5B09B8AD

*Locusta migratoria*

urn:lsid:zoobank.org:pub:D792A09E-844A-412A-BFCA-5293F8388F8C

*Periplaneta americana (Blatta americana)*:

urn:lsid:zoobank.org:act:95113A55-4C6D-4DC7-A0E5-620BACADFFE5

*Apis mellifera*:

urn:lsid:zoobank.org:act:9082C709-6347-4768-A0DC-27DC44400CB2

*Bombyx mori (Phalæna (Bombyx) mori)*

urn:lsid:zoobank.org:act:215466E3-E77F-46E9-8097-372837D7A375

*Drosophila melanogaster*:

urn:lsid:zoobank.org:act:5B39F0AA-270D-4AA8-B9A3-C36A3A265910

## Introduction

Like in other organisms, pathogens seem to be important drivers of evolution in social insects. On the one hand, social insects are a ‘desirable’ target for pathogens as an insect colony represents a large source of many potential hosts, all with a similar genetic background. Thus, if pathogens manage to enter a colony they can exploit many individuals. However, social insects are also well protected as they evolved ‘social immunity’ (Cremer et al., 2007), a repertoire of defensive mechanisms that work at the colony level. For instance, behavioral task division limits contact to potentially infected individuals and can lead to their eviction. Also molecular mechanisms of social immunity exist, for example indirect immunization of colony members (e.g., Cremer et al., 2007; Masri and Cremer, 2014) or impregnation of the nest walls with fungicidal compounds (Rosengaus et al., 2011). Thus, social immunity can be considered a selected emergent property of insect colonies where the whole is more than the sum of the individuals. Social immunity is an evolved social trait that aligns with the complexity of social organization. So the question might arise, can pathogens even be a driver of complex social organization and does this vary across different social species with different ecologies?

In termites, there is evidence that implies that pathogen load may have been an important driver of brood care when comparing species of early branching lineages of the termite phylogeny (Korb et al., 2012). These species share a similar life type of nesting in a single piece of wood that serves as food and shelter (one-pieces nesting or wood-dwelling termites, see Abe, 1987; Korb, 2007). ‘Workers’ are developmentally totipotent immatures that can become winged as well as wingless reproductives. Brood care by workers is less developed in these wood-dwelling species where all individuals sit inside their food. Interactions are generally not altruistic but cooperative as all individuals function equally as donors and recipients (Korb, 2007). Yet, the degree of brood care seems to be malleable depending on the pathogen load. It is negligible / absent in several drywood termites (Kalotermitidae) that nest in sound dry wood with a low pathogen load (Rosengaus et al., 2003). However, the dampwood termite (Archotermopsidae), *Zootermopsis nevadensis*, which nests in rotten, decaying wood with many fungi and other potential pathogens, show more intensive brood care mainly consisting of allogrooming (Rosengaus et al., 1999; Korb et al., 2012). However, even in the drywood termites, parasites can affect social behavior. Infestations of *Cryptotermes secundus* colonies with mites alter ‘reproductive decision making’ as workers of infested colonies (even if they are not themselves infected) develop into winged sexuals that leave the nest during nuptial flights (Korb and Fuchs, 2006). In the dampwood termite, *Z. nevadensis*, pathogens present a permanent threat which is reflected in huge constitutive investments in immune defense at different levels, the individual as well as the colony level, potentially with a socially acquired immunity (Rosengaus et al., 1999; Traniello et al., 2002; Bulmer et al., 2009; Rosengaus et al., 2011). This is also reflected in *Z. nevadensis*’ genome where six copies of Gram-negative binding proteins (GNBPs) were found, more than in any other insect species (Terrapon et al., 2014). Four of these genes were also found in the genome of the fungus growing termite *Macrotermes natalensis* (Termitidae) and they seem to be termite specific (Poulsen et al., 2014; Korb et al., 2015). *M. natalensis* is not a wood-dwelling species but a foraging species with intensive brood care and complex social organization. It builds the nest (mound) separate from the foraging grounds so that individuals leave the nest to forage and bring back food to the central nest (i.e., central place foraging). Thus, it also is constantly exposed to pathogens, but in contrast to wood-dwelling dampwood termites, it encounters pathogens mainly during foraging. For other central-place foraging termites, such as several Australian *Nasutitermes* species (Termitidae) which have a complex social organization., GNBPs had previously been shown to be under positive selection (Bulmer and Crozier, 2004; Bulmer and Crozier, 2006; Bulmer et al., 2010).

Based on these results, the hypothesis was proposed that selective pressure on immune defense genes (IGs) differs across termites depending on their life type (Korb et al., 2015). We tested the hypothesis that wood-dwelling termites, which do not leave the nest to forage outside show relaxed selection on immune genes and fewer signs of positive selection compared to soil-foraging species. Additionally, we tested whether within the wood-dwelling life type, the dampwood termite *Z. nevadensis* had stronger signs of selection than other wood-dwellers that nest in sound wood.

In order to test for differences in selective forces acting on IGs between wood-dwelling and foraging species, we analyzed a set of 81 previously identified single-copy IGs (see Materials and Methods) in eight termite species (published data see Misof et al., 2014; Terrapon et al., 2014; Poulson et al., 2014; Harrison et al., 2018; Evangelista et al., 2019; data sources are provided in Supplementary Table S1). Four of the species we investigated are wood-dwelling species: *C. secundus, Incisitermes marginipennis, Prorhinotermes simplex*, and *Z. nevadensis*. The remaining four species are foraging species: *Mastotermes darwiniensis, Reticulitermes santonensis* (i.e. *R. flavipes*), *Coptotermes* sp. And *M. natalensis*. These were analyzed in a phylogenetic framework of 22 species (one mayfly, twelve polyneopteran insects including above listed termites and their closest relatives Cryptocercidae, two paraneopteran and six holometabolous insects) spanning winged insects to increase the statistical power of lineage specific tests for selection.

## Results

### Phylogenetic relationships

The species tree was inferred from a supermatrix including 1178 single-copy orthologs (SCOs) and spanning an alignment length of 555,906 amino acid positions (partition coverage 100%, site-completeness score C_a_ = 74.61%, see Supplementary Materials and Supplementary Figure S1). Termites were monophyletic with *Cryptocercus* as sister group, consistent with earlier work (e.g., Lo, 2000; Klass, 2006; Inward et al., 2007; Legendre et al., 2008). Phylogenetic relationships within termites are largely consistent with Evangelista et al. (2019) and are statistically maximally supported (Figure 1). Consistent with earlier work (e.g., Legendre et al., 2008), neither wood-dwellers nor foragers constitute monophyletic groups, confirming that several independent switches in life type were included in our analyses. More details on phylogenetic analyses are provided in the Supplementary Materials.

**Figure 1.**
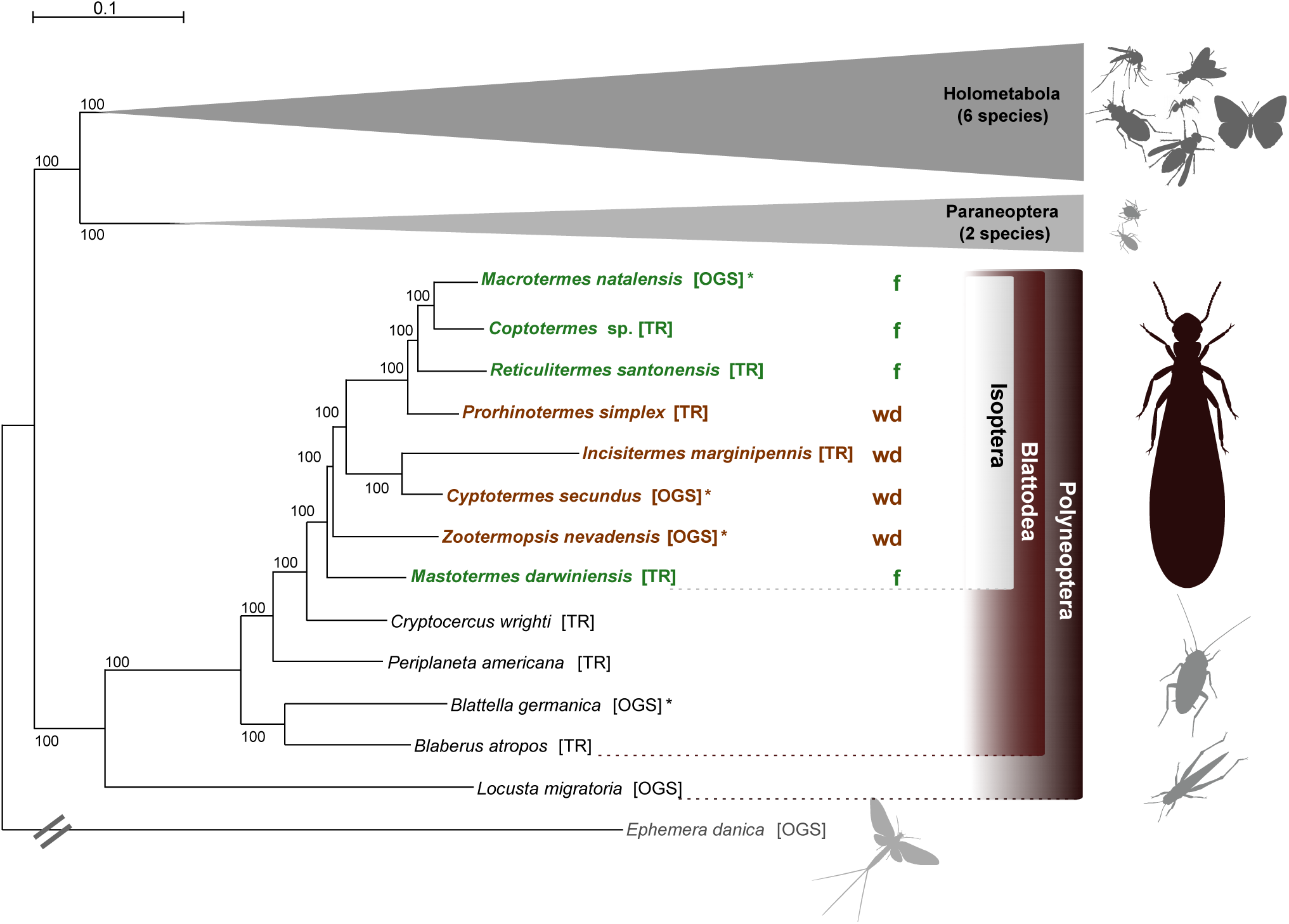
Phylogenetic relationships. Best ML species tree (out of 50 trees) inferred from a superalignment of 1178 single-copy orthologs (SCOs) with 555,906 amino acid positions (schematized for Holometabola and Paraneoptera, all ML trees had the same topology). Statistical bootstrap support was derived from 200 replicates. The tree was rooted with the mayfly *Ephemera danica* (the full tree is provided with the Supplementary Files on DRYAD). Color code: brown indicates wood-dwelling species (wd), green indicates foraging species (f). TR: derived from transcriptome assemblies (Misof et al., 2014; Evangelista et al., 2019); OGS: derived from official gene sets (Supplementary Table S1). * indicates reference species whose OGS was used to create the ortholog set. Pictograms were kindly provided by H. Pohl, Jena, (adopted from Misof et al., 2014).

### Patterns of selection on termite immune genes

Between 13 and 78 single-copy orthologs of IGs per species were included in the analyses (Table 1). We found no evidence for positive selection on the IGs (see Supplementary Table S2).

**Table 1.**
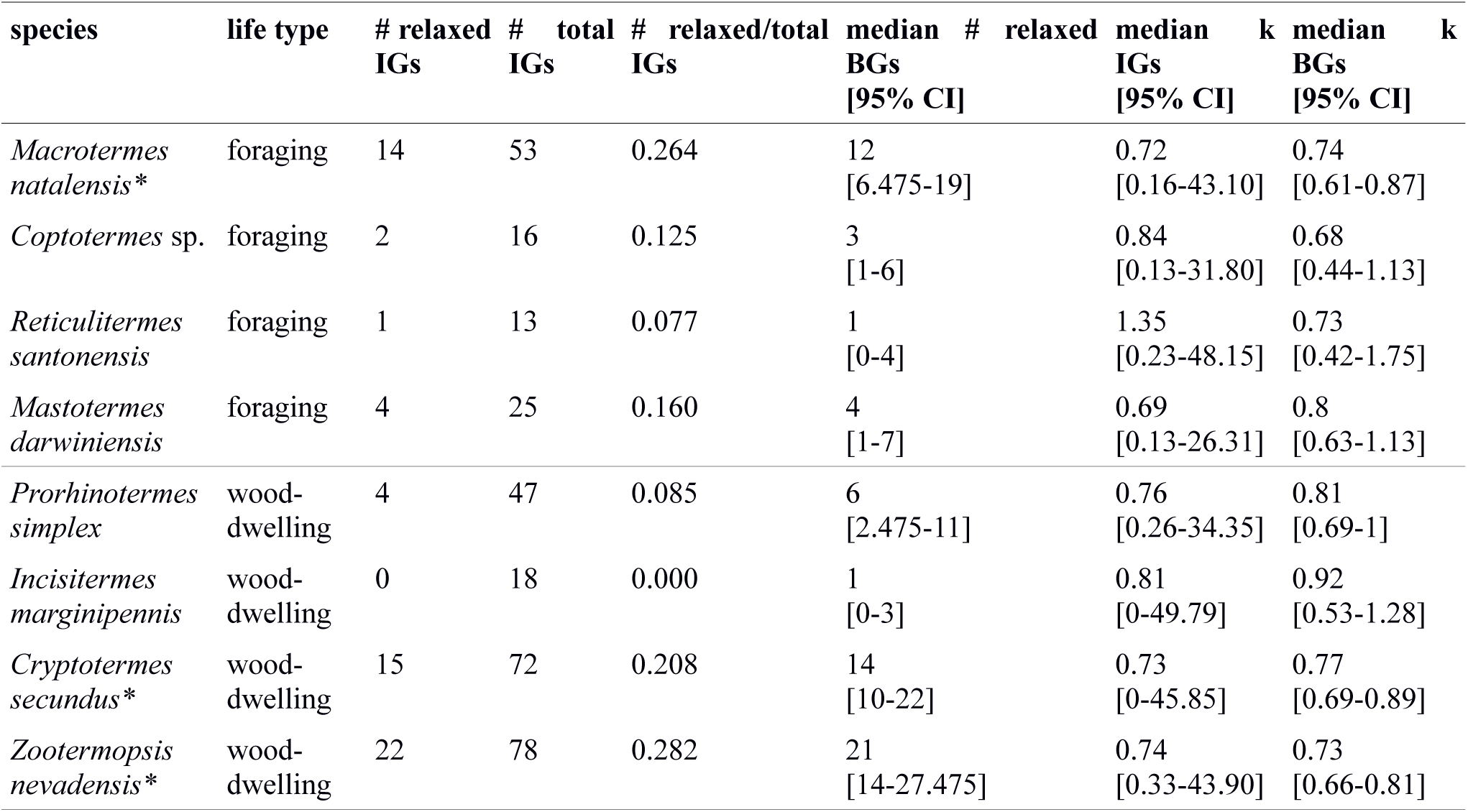
Results of RELAX analyses of changes in selection intensity on termite immune genes (IGs). Genes that are under significantly relaxed selection were counted (k < 1, *P* < 0.05) for columns containing #. BG: background single-copy ortholog. * indicates species with annotated genomes.

Next, we tested the hypothesis that selection on the IGs of wood-dwelling species is relaxed compared to foragers. We found 62 cases of significantly relaxed selection across all termite species analyzed (Table 1, Supplementary Table S3). There was no evidence for a difference in the number of IGs under relaxed selection between the life types (generalized linear mixed effects model with binomial error distribution: df = 7, z = −0.191, *P* = 0.85). There was also no evidence for differences between species (generalized linear model assuming binomial error distribution: df = 7, z = −1.74 – 0.729, *P* = 0.082-1, ranges of z and *P* are for the different species). Because changes in the selection intensity on IGs could be obscured by genome wide differences in selective constraint, it is important to test these hypotheses against the genomic background. To this end, we generated sets of background genes (BGs) that consisted of genes matching the GC-content and sequence length of each IG for each species (see Materials and Methods). The number of IGs under relaxed selection did not differ significantly from that expected from the analysis of the BGs for any of the species (see 95% confidence intervals (CIs) for BG sets in Table 1). In order to take genome-wide effects of selective constraint into account, when comparing selection intensity between wood-dwellers and foragers, we (i) calculated the ratio of genes under significantly relaxed selection between wood-dwellers and foragers for IGs and (ii) compared this ratio to that expected from BGs (see Materials and Methods). If selection is relaxed specifically in the IGs of wood-dwellers relative to their genomic background, we expect the ratio of the number of significant genes under relaxed selection between wood-dwellers and foragers to be larger for the IGs than for the BGs. However, the ratio of the number of IGs under relaxed selection between wood-dwellers and foragers did not differ significantly from the expectation derived from BGs (Figure 2A), supporting the view that patterns of relaxed selection on IGs follow genome wide trends in selective constraint.

**Figure 2.**
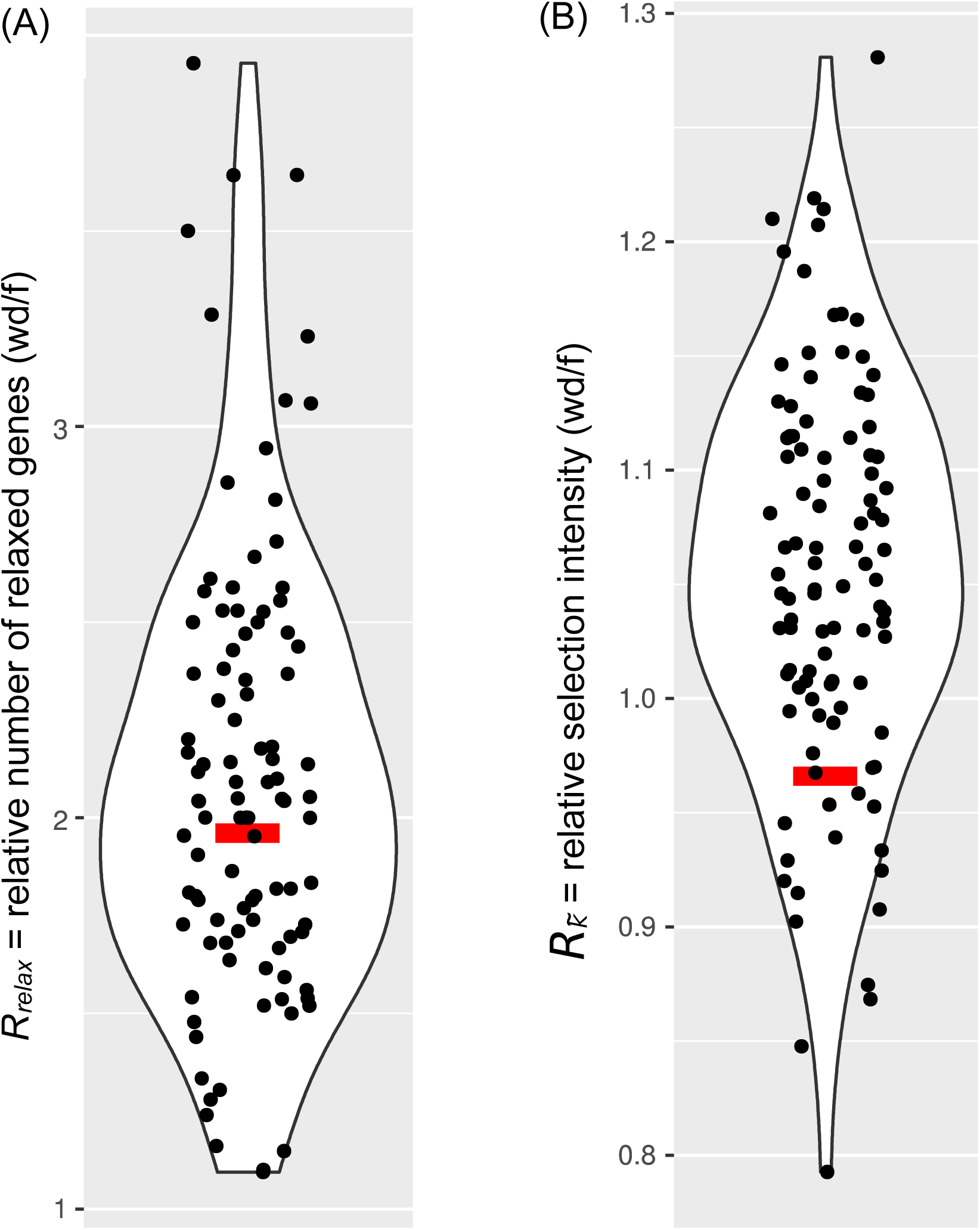
Comparison of selection on immune genes to sets of genes that represent the genomic background. Relaxed selection on IGs does not differ from the genomic background. Red bar = immune genes (IGs); black dots = 100 sets of background genes (BGs). (A) The ratio of the number of genes under significantly relaxed selection between wood-dwellers and foragers (*R*_*relax*_). (B) The relative intensity of selection (*R*_*k*_) as measured by the ratio of median k between wood-dwellers and foragers.

Because we did not find any evidence for an increase in the number of IGs under significantly relaxed selection in wood-dwellers, we reasoned that a putative signal of relaxation of selective constraint might be more diffuse and only become visible as a general trend over all IGs investigated. In order to capture such more general trends, we assessed potential differences in k, a measure for the intensity of selection, for all IGs between life types. K did not differ significantly between species (Kruskal-Wallis test: df = 7, χ^2^ = 3.95, n = 322, *P* = 0.79, for n per species see Table 1) nor was it lower for wood-dwellers (Mann-Whitney U test, one-sided: U = 11690, n = 322, *P* = 0.41). This indicated similar selection intensity on IGs for wood-dwellers and foragers. For the comparison of k between life types it is, as above, important to take the selective constraint on the genomic background into account. K for the IGs did not differ significantly from k for sets of BGs for any of the species investigated (Table 1). Following the same rationale as above, we used the ratio of medians of k between wood-dwellers and foragers as a test statistic. This ratio can be interpreted as the relative intensity of selection between wood-dwellers and foragers. We found that the relative intensity of selection between wood-dwellers and foragers on IGs matched that of the BG sets (Figure 2B), again suggesting that the IGs follow genome-wide trends of selective constraint.

Finally, we hypothesized that our results might be affected by the particular selection pressures that act on *Z. nevadensis*, which is a wood-dwelling species, but lives in relatively microbe-rich damp wood (Rosengaus, 2003). Hence, selection would be expected to be stronger on *Z. nevadensis* IGs than on IGs in the other wood-dwellers, resulting in a smaller fraction of genes under significantly relaxed selection in *Z. nevadensis*. Contrary to this expectation, we found significantly more IGs under relaxed selection in *Z. nevadensis* (generalized linear model assuming binomial error distribution: df = 3, z = 2.53, *P* = 0.01). However, when we excluded *Z. nevadensis* from the comparison between life types, there was also no difference in the number of IGs under relaxed selection (generalized linear mixed effects model with binomial error distribution: n = 8, z = −0.94, *P* = 0.35), showing that our results are robust. The overall intensity of selection on IGs (k) did not differ significantly between *Z. nevadensis* and the other wood-dwellers (Mann-Whitney U test, U = 5471, n = 215, *P* = 0.77). Similarly, it could be argued that the assumption of relaxed selection only holds for the dry wood-dwellers *C. secundus* and *I. marginipennis*, assuming that only dry wood is a truly pathogen poor substrate. We could not find a difference in the number of genes under relaxed selection between the dry wood-dwellers and the other species (generalized linear model with binomial error distribution: df = 7, z = 0.73, *P* = 0.464) nor for the intensity of selection over all IGs as measured by k (Mann-Whitney U test: U = 10538, n = 322, *P* = 0.37).

To our surprise, we observed that median k for IGs was smaller than one for seven of the eight investigated termite species (Table 1). This difference is significant (Mann-Whitney U test: U = 20958, n = 322, *P* < 0.01), indicating an overall relaxation of selection. This signal was not IG specific: k for the sets of BGs was also significantly smaller than 1 for all species (P = 4×10e-18 - 4.8×10e-5), suggesting a genome-wide relaxation of selection on the termite lineages compared to the background branches of the phylogeny.

## Discussion

In this study we combined recent genomic and transcriptomic resources to test whether termite ecology, in particular exposure to pathogens, might affect the evolution of immune genes. Surprisingly, and in contrast to studies from *Drosophila* (Clark et al., 2007; Sackton et al. 2007; Hill et al., 2019) we could not find evidence for positive selection on the immune genes for eight termite species. We extended our analyses, employing recent tests to explicitly assess relaxed selection (Wertheim et al., 2015), revealing 62 cases of significantly relaxed selection in IGs.

We expected that the intensity of selection would differ between termites of the wood-dwelling and foraging life types, as foraging termites are assumed to experience higher selection pressure from pathogens due to higher exposure. Contrary to our expectation, we did not detect an effect of life type on signs of selection for immune genes. Neither did wood-dwelling species differ from the soil foraging species, nor did the dampwood termite *Z. nevadensis* differ from the other wood-dwelling termites. *Z. nevadensis* that has a higher microbial load compared to the other wood-dwellers (Rosengaus, 2003), even had an elevated number of IGs, for which selection was significantly relaxed. However, the number of IGs under relaxed selection in *Z. nevadensis* did not differ from the expectation derived from BGs, suggesting that this is not an IG specific signal. Taken together, this suggests that individual immune response remains an important trait in termites even if social immunity is effective or pathogen exposure is low to moderate. A possible explanation for similar selection pressure on immune genes between wood-dwelling and foraging species is that both harbor complex gut microbial communities that are essential for termite survival (see Waidele et al., 2017). It seems reasonable to assume that the immune system is likely to play a role in modulating these communities in wood-dwellers and foragers alike resulting in constant selection pressure.

We applied state of the art methods (Pond et al., 2005; Murrell et al., 2015; Wertheim et al., 2015) and carefully controlled our study by analyzing background genes that matched IGs in GC-content and sequence length (see Materials and Methods). We also combined, to our knowledge, the largest data set of termite genomic and transcriptomic resources for our study that has been used in that context so far. Nonetheless, our study might lack statistical power: First, we have relatively few species for life type comparisons (4 vs 4). Second, for five of these species (*M. darwiniensis, I. marginipennis, R. santonensis, P. simulans* and *Coptotermes* sp.) only transcriptome data are available. Naturally, the number of genes under significantly relaxed selection is lower for the species for which only transcriptome data are available, as fewer genes could be annotated and investigated in the transcriptome data (Table 1, 2 and Supplementary Table S4). However, potential biases in data availability were taken into account in the procedure to sample sets of matching background genes.

**Table 2.**
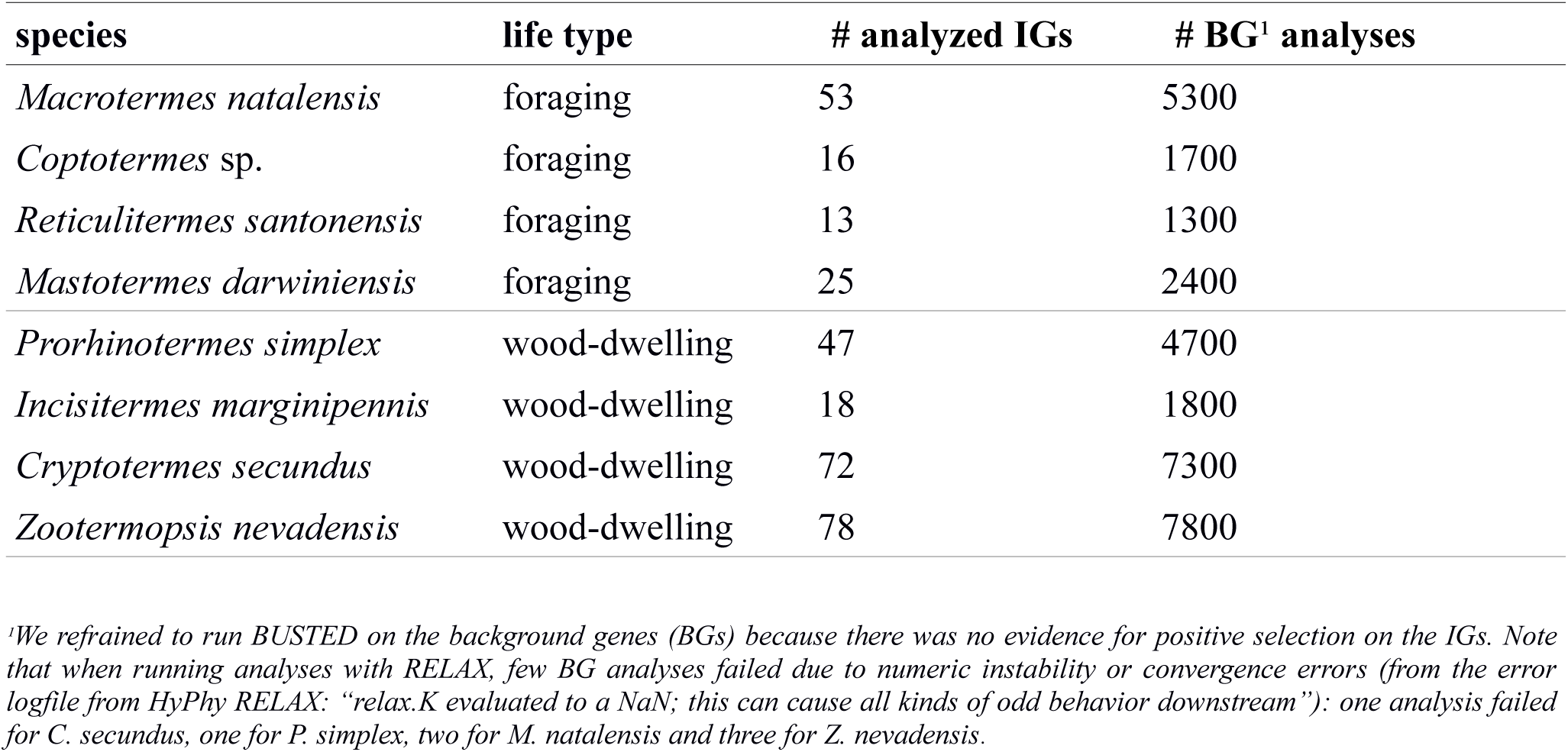
Number of analyzed IGs with HyPhy-BUSTED and RELAX and respective BG analyses with HyPhy-RELAX for each of the included termite species.

| species | life type | # analyzed IGs | # BG<sup>1</sup> analyses |
| --- | --- | --- | --- |
| *Macrotermes natalensis* | foraging | 53 | 5300 |
| *Coptotermes* sp. | foraging | 16 | 1700 |
| *Reticulitermes santonensis* | foraging | 13 | 1300 |
| *Mastotermes darwiniensis* | foraging | 25 | 2400 |
| *Prorhinotermes simplex* | wood-dwelling | 47 | 4700 |
| *Incisitermes marginipennis* | wood-dwelling | 18 | 1800 |
| *Cryptotermes secundus* | wood-dwelling | 72 | 7300 |
| *Zootermopsis nevadensis* | wood-dwelling | 78 | 7800 |

We refrained to run BUSTED on the background genes (BGs) because there was no evidence for positive selection on the IGs. Note that when running analyses with RELAX, few BG analyses failed due to numeric instability or convergence errors (from the error logfile from HyPhy RELAX: “relax.K evaluated to a NaN; this can cause all kinds of odd behavior downstream”): one analysis failed for *C. secundus*, one for *P. simplex*, two for *M. natalensis* and three for *Z. nevadensis*.

When taking the genomic background and the sampling procedure into account (Figure 2), our data provide no evidence that selection intensity on single-copy ortholog IGs differs between life types. Do our results imply that selection pressure on immune genes does not differ between termite life types? No, we cannot conclude this as our study excluded immune genes that are not single-copy orthologs. Antimicrobial peptides (AMPs) or Gram-negative binding proteins (GNBPs) that have several copies within the genome of some species were not addressed in our study. Different copies can be caste-specifically expressed in termites, as shown for *Z. nevadensis (*Terrapon et al., 2014), and for *Reticulitermes speratus* (Mitaka et al., 2017), and they may be under positive selection as indicated, for instance, by studies on GNBPs for several foraging *Nasutitermes* species (Bulmer and Crozier, 2006; Bulmer et al., 2010). Note that the original hypothesis for a difference in immune defense between wood-dwellers and foraging termites considered specifically GNBPs and AMPs (Korb et al., 2015). Thus, more studies are warranted that test selection on such multi-copy genes. However, orthology is difficult to infer for multi-copy genes making the restriction to SCOs a standard procedure (e.g., Pauli et al., 2016; Dowling et al., 2016; Mitterboeck et al., 2017; Ran et al., 2018; Brandt et al., 2019; Hill et al., 2019). Thus, we restricted our analysis to single-copy orthologs because orthology is a basic assumption of the current methods that identify selection in a powerful phylogenetic framework (e.g. codeML implemented in PAML, Yang, 2007; HyPhy applying BUSTED, see Murrell et al., 2015, and RELAX, see Wertheim et al., 2015).

Applying state of the art methods on the most comprehensive termite data set for which genome or transcriptome data are currently available, we found no evidence for differences in selection on immune genes that correlate with termite ecology, specifically pathogen exposure. Our results suggest that the putative evolutionary response to differences in pathogen exposure can not be found in single-copy immune genes. Interestingly, we detected a signal of overall relaxation of selective constraint in termites. This could be related to smaller effective population sizes due to their social organization opening up an interesting new line of research (Rubinstein et al., 2019).

## Materials and Methods

### Identifying orthologous sequence groups of protein-coding single-copy genes

As basis to identify single-copy orthologs (SCOs), we designed an ortholog set from official gene sets (OGS) from available full (draft) genomes of four species: *Cryptotermes secundus, Macrotermes natalensis, Zootermopsis nevadensis*, and *Blattella germanica* (Supplementary Table 1). The set of single-copy orthologs was inferred with the software OrthoFinder v.1.1.4 (Emms and Kelly, 2015) using default settings. As input, the OGS of respective species were downloaded from public databases as amino acid and nucleotide sequences. The OGS of *C. secundus* was kindly provided by the *C. secundus* consortium (via J. Korb) before it was published (Harrison et al., 2018). We only kept the longest isoform per orthologous group (OG). All OGs that included the amino-acid Selenocysteine (U) were removed since they might be putative pseudo-genes and to avoid difficulties in downstream analyses as many software packages are not able to handle Selenocysteine. This was done using the package Biobundle (script isoformCleaner with boost 1.61.0 environment (Kemena, 2017, available from https://github.com/CarstenK/BioBundle). SCOs inferred with OrthoFinder, were summarized with custom-made Python scripts (kindly provided by A. Faddeeva and L. Wissler, available upon request). This resulted in a set of 5382 SCOs across the four selected species.

### Taxon sampling

We included the four reference species that were used to create the ortholog set as well as genome and transcriptome data of 18 additional species in our analyses. Eight of the included species are termites: *Coptotermes* sp., *Incisitermes marginipennis, Mastotermes darwiniensis, Prorhinotermes simplex*, and *Reticulitermes santonensis* (with published transcriptome data); *C. secundus, Z. nevadensis* and *M. natalensis* (with published OGS), see Evangelista et al. (2019). Other included species were a representative of Cryptocercidae as it is supposed to be the sister group of termites (e.g., Lo et al., 2000; Inwards et al, 2007), two other non-social cockroach species, and representatives from other polyneopteran, paraneopteran and holometabolous insects, and a mayfly as outgroup (Adams et al., 2000; Bonasio et al., 2010; Elsik et al., 2014; Evangelista et al., 2019; Harrison et al., 2018; International Aphid Genomics Consortium, 2010; Misof et al., 2014; Mesquita et al., 2015; Pauli et al., 2016; Poulsen et al., 2014; Richards et al., 2008; Sinkins, 2007; Terrapon et al., 2014; Wang et al., 2014; Xia et al., 2004; the full species list is provided in Supplementary Table S1). Access to transcriptome data (see Figure 1) was kindly granted by 1KITE before they were published, access to the OGS of the locust and the mayfly was granted by the i5K community.

### Assignment of putative orthologous transcripts to the SCOs

The ortholog set was used as input for the assignment of putative SCOs (provided as Supplementary files on DRYAD). Inference and assignment of putative orthologs from genome and transcriptome data of the 18 species that were not included for generating the ortholog set was performed with Orthograph v.0.6.1 (Petersen et al., 2017). Orthograph is recommended to infer orthologs from transcriptome data for which no official gene set are available (see Petersen et al., 2017). Orthograph analyses resulted in 5366 SCOs that were identified in at least one species that was not used as reference species to create the ortholog set.

### Multiple sequence alignments, species tree inference and testing for selection

Individual SCOs were aligned at the amino acid level with MAFFT v7.310 using the L-INS-i algorithm (Katoh and Standley, 2013).

### Species tree inference

For inferring the species tree, we only kept those SCOs that were present in all 22 species. This resulted in 1178 SCOs. Ambiguously aligned sections on the amino acid level were identified with Aliscore v2.2 (Misof and Misof, 2009; Kück et al., 2010) (settings: -r with all pairwise sequence comparisons, -e for gap-rich alignments, otherwise defaults) and masked with Alicut v2.3 (Kück, 2011). Masked amino-acid multiple sequence alignments (MSAs) were concatenated into a supermatrix with FASconCAT-G v.1.02 (Kück and Longo, 2014). We inferred phylogenetic relationships using a maximum-likelihood (ML) approach with IQtree v1.5.4 (Nguyen et al., 2014; Chernomor et al., 2016). Statistical support was determined from 200 non-parametric, slow and thorough bootstrap replicates. We ensured bootstrap convergence with *a posteriori* bootstrap criteria (Pattengale et al., 2010) as implemented in RAxML (Stamatakis, 2014), v.8.2.11. The best ML tree, out of 50 inferred trees, which all showed an identical topology, was rooted with *Ephemera danica* using SeaView v.4.5.4 (Gouy et al., 2010; note that multiple tree viewers are not reliable, see Czech et al., 2017); trees were graphically edited with Inkscape (v.0.91) (www.inkscape.org). More details on the procedure of phylogenetic inference are provided in the Supplementary Materials.

### Inferring natural selection

### Alignment processing and clean-up

Methods to identify selection are sensitive to misalignments (Privman et al., 2012; Markova-Raina and Petrov, 2011). Therefore we performed extensive alignment cleanup. First, we identified and deleted badly aligned or gap-rich sequences on amino acid level with MaxAlign v1.1 (Gouveia-Oliviera et al., 2007). This procedure resulted in five SCOs with only one sequence which were excluded from further analyses. We subsequently compiled corresponding nucleotide (i.e, codon) MSAs with PAL2NAL (Suyama et al., 2006, v14.1, see Misof et al., 2014) using the 5361 amino-acid MSAs as blue-print. The nucleotide MSAs were then used for all following analyses. Second, we deleted all SCOs with less than four sequences (223 SCOs) leaving 5138 SCOs. Third, we identified ambiguously aligned sections on amino acid level with Aliscore v2.2 (Misof and Misof, 2009; Kück et al., 2010) with the same settings as described for the species tree inference. Suggested sections were removed from the amino acid and correspondingly from the nucleotide MSAs with Alicut v2.3 (Kück, 2011). Subsequent analyses were performed on the masked nucleotide MSAs. First, we classified 5138 SCOs into 86 immune single-copy genes (IGs) and into the remaining 5052 SCOs based on Terrapon et al. (2014, see Table S25 for *Z. nevadensis*) and Korb et al. (2015, for *Z. nevadensis* and *M. natalensis*), see Supplementary Table S5. The 5052 non-immune SCOs were used to generate gene sets from the genomic background, i.e. background genes (BGs) that had similar GC-content and sequence length (see below) as the examined IGs. Note that from the 86 IGs (Supplementary Table S5) five IGs were excluded because there was no SCO fulfilling the criteria to serve as BG and these were not listed by Terrapon et al. (2014) or not reported by Korb et al. (2015). This left 81 IGs for analyses (for detailed information see Supplementary Materials).

To further reduce potential false positives that may originate from misalignments, we trimmed trailing ends of each multiple sequence alignment, i.e. each multiple sequence alignment started and ended with unambiguous nucleotides for all species. Because visual inspection of the trimmed MSAs still revealed putative misaligned nucleotides, we applied the GUIDe tree based AligNment ConfidencE approach (GUIDANCE) Guidance2 (Landan and Graur, 2008; Sela et al., 2015) version 2.02 using MAFFT as implemented alignment method on the trimmed MSAs (options: codon as sequence type, sequence cutoff=0 and the default column cutoff=0.93).

### Inferring positive selection and selection intensity

To test for evidence of positive selection we used BUSTED (Murrell et al., 2015) as implemented in the software package HyPhy (Pond et al., 2005). BUSTED uses a branch-site test for positive selection on entire genes in a foreground branch relative to the background branches in a phylogeny. A significant *P*-value means that at least one codon in the foreground branch has experienced an episode of positive selection. The high sensitivity of the method compared to tests from alternative packages (see e.g., Enard et al., 2016; Ebel et al., 2017; Venkat et al., 2018; Hill et al., 2019) and the option to define the foreground branches according to our research question made it perfectly suited for our study.

For inferring potential relaxation of selection, we used RELAX (Wertheim et al., 2015) as implemented in the software package HyPhy (Pond et al., 2005). RELAX has been designed to identify changes in the intensity of selection on a given protein-coding gene in a codon-based phylogenetic framework (see Wertheim et al., 2015). The basic expectation of RELAX is that under relaxed selection, the ω of sites under purifying and positive selection will move closer to neutrality. The change of ω for the selected sites relative to the background branches is quantified with the selection intensity parameter k, where

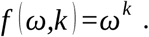

If parameter k is significantly larger than one, selection has been intensified along the test branches. If k is significantly smaller than one, selection has been relaxed.

We used BUSTED and RELAX as implemented in the software package HyPhy, version 2.4.0-alpha.2 (access: April 2019). We determined k for each of the termite species relative to all other species in the tree. We chose this species-wise analysis setup because we wanted to take species level differences in pathogen exposure into consideration.

### Comparison of IG selection parameter to the genomic background

To test whether or not signals of selection were specific to IGs or the consequence of genome-wide trends, we generated sets of background genes (BGs). To this end, we searched the 5052 non-immune SCOs for genes that closely matched the GC-content (+/− 5%) and the sequence length (+/− 5%). Following this procedure, we generated lists of matching BGs for each IG and each species. From these lists, we randomly sampled 100 gene sets such that there was a matching BG for each IG that was analyzed in the respective species (e.g., for *Z. nevadensis*, 78 IGs were analyzed, thus each of the 100 sets of BGs contained therefore 78 BGs, see also Table 2. Lists of analyses BG that are similar in GC-content and sequence length of the IGs are provided for each species as Supplementary files on DRYAD). We performed BUSTED and RELAX analyses for each termite species on all IGs and RELAX on the species’ respective BG set with default settings.

All analyses were performed on Linux Desktop PCs at the University of Freiburg, Germany and on the Linux HPC CSIRO Cluster Pearcey, Australia. Analyses results of all IGs are summarized in Supplementary Table S2, S3; results of BGs are provided species-wise on DRYAD.

### Statistical analyses

In order to assess potential differences in selection intensity on the IGs between species and life types, we summarized the RELAX results by counting the number of genes under significantly relaxed selection: genes with k < 1 and *P* < 0.05 were considered. Potential differences were tested for statistical significance with generalized linear models with binomial error distribution using the functions glm and glmer from the lme4 R package (Bates et al., 2015, version 1.1-21 with R (2018), version 3.4.4). The number of significant genes divided by the total number of genes analyzed was used as response variable. Species or life type were used as potential predictors. Varying sampling depths between species, as represented by the number of IGs analyzed per species, were taken into account as weights in the model. When comparing life types, species were treated as a random effect. See Supplementary file (RanalysisscriptfortermiteIGs.R on DRYAD) for a detailed R analysis script with all models, commands and functions used.

We also analyzed parameter k to search for more diffuse trends in selection intensity that are distributed over the IGs so that individual IGs do not reach significance. According to its definition, k should map linearly on a logarithmic scale. However, we found six strong outliers on the logarithmic scale that were more than standard deviations away from the mean (log(k) < −9, see Supplementary Table S3) that could make the analysis in a linear framework error prone. Visual inspection of the alignments underlying these extremely small values of k did not reveal any obvious misalignments that would justify their exclusion. Therefore, potential differences in k were assessed with non-parametric tests (Mann-Whitney U test, Kruskal-Wallis test) that are robust to outliers.

Genome-wide trends in selection intensity can potentially obscure IG specific patterns or generate false positives. For example, changes in population size can affect the efficiency of both purifying and positive selection (Ohta, 1973) on a genome wide scale. Population sizes might differ between species and life types in our study depending on reproductive rates and degrees of sociality. Therefore, it is essential to put the results for IGs into the context of the genomic background. To this end, we generated expected values for the number of significant genes and for k based on 100 sets of BGs (see above) per termite species, representing the genomic background. The median of k and 95% CIs from the BG-based distributions for each species were calculated with R (version 3.4.4), using the median and quantile functions with standard settings. In order to compare differences between life types while taking the genomic background into account, we calculated (i) the ratio of the number of genes that were significantly relaxed between wood dwellers and foragers

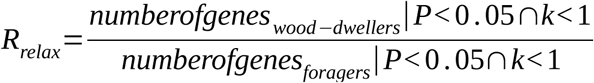

and (ii) the ratio of median parameter k (i.e., relative selection intensity):

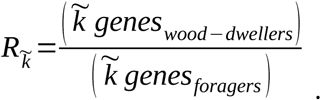

These ratios were calculated for the IGs and the BGs. Then the ratio of the IGs was compared to their expectation from the BGs. Significant shifts in selection intensity that are specific to IGs should lead to shifts of *R*_*relax*_ and 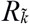 only for IGs. Thus, if there were IG specific patterns of relaxed selection, the ratios *R*_*relax*_ and 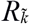 for the IGs should represent extremes of the distribution of sets of BGs.

## Data Availability Statement

Supplementary data (see Supplementary Materials) used in this study can be found at the DRYAD digital repository, DOI XXX. Will be available upon manuscript acceptance.

## Author Contributions

JK and FS conceived the study; KM, MS, FS performed all analyses; FS, KM and JK wrote the manuscript. All authors edited and approved the Manuscript.

## Acknowledgments

We acknowledge Bernhard Misof, Panos Provataris, Coby Schal, Xavier Belles, Stephen Richards and the i5K community the usage of the official gene set of *Ephemera danica*, and *Blattella germanica*, and Xianhui Wang and Le Kang for access and usage of the official gene set of *Locusta migratoria*. We also thank the 1KITE Dictyoptera group who granted us access to transcriptome assemblies before they were published. We thank David Enard (University of Arizona, USA), Ondrej Hlinka (CSIRO, Australia), Ryan Velazquez and Sergej Pond for their useful input and help with analyses of the molecular sequence data using Guidance2 and HyPhy. KM thanks Thomas Pauli (University of Freiburg, Germany) for fruitful discussions and Hans Pohl (University of Jena, Germany), who kindly allowed the usage of several pictograms in Figure 1 (adopted from Misof et al., 2014).

## Supplementary Materials

The Supplementary Materials for this article can be found online at:

